# Boundary effects of expectation in human pain perception

**DOI:** 10.1101/467738

**Authors:** E.J. Hird, C. Charalambous, W. El-Deredy, A.K. Jones, D. Talmi

**Affiliations:** University of Manchester, Manchester, UK; Institute of Psychiatry, Psychology and Neuroscience, Kings College London, London, UK; Centro de Investigación y Desarrollo en Ingeniería en Salud, Universidad de Valparaiso, Chile; Salford Royal NHS Foundation Trust, Manchester, UK

## Abstract

Perception of sensory stimulation is influenced by numerous psychological variables. One example is placebo analgesia, where expecting low pain causes a painful stimulus to feel less painful. Yet, because pain evolved to signal threats to survival, it should be maladaptive for highly-erroneous expectations to yield unrealistic pain experiences. Therefore, we hypothesised that a cue followed by a highly discrepant stimulus intensity, which generates a large prediction error, will have a weaker influence on the perception of that stimulus. To test this hypothesis we collected two independent pain-cueing datasets. The second dataset and the analysis plan were preregistered (<u>osf.io/5r6z7</u>). Regression modelling revealed that reported pain intensities were best explained by a quartic polynomial model of the prediction error. The results indicated that the influence of cues on perceived pain decreased when stimulus intensity was very different from expectations, suggesting that prediction error size has an immediate functional role in pain perception.

## Introduction

The experience of pain results from both sensory input and psychological variables such as personality traits and anxiety level ^1–5^. One important psychological variable is how intense one expects that imminent pain might feel. Expectations about a painful event shift the perceived intensity of pain closer to the expected intensity. For example, in placebo analgesia, expecting low pain decreases pain perception and associated brain activity ^6–11^. Likewise, expecting high pain increases the perceived intensity of pain, as in nocebo hyperalgesia ^12^.

Imagine having a calm picnic in the garden, when you are suddenly stung by a wasp. Although you had no reason to expect any pain before being stung, you immediately feel searing pain. The discrepancy between prior pain expectations, which in this example were null, and the sensory evidence, here the bodily response to the venom in the sting, is termed pain prediction error (PE). It is not clear whether the size of that prediction error has any immediate functional role in how pain is perceived. It seems clearly adaptive for expectations to modulate sensory perception to some extent ^13, 14^. Available evidence indicates that experiencing large PEs leads to learning over time ^15, 16^. However, failing to adjust perception based on the immediate sensory reality could lead to inaccurate, possibly hallucinatory perception, as described in recent theories of psychosis ^17^. Therefore, when prediction error is high, it may make sense for perception to be influenced more strongly by the sensory input. In the example of the wasp sting, the most adaptive response would be for your prior expectation: “calm picnic” to have less impact on pain perception than the sensory input: “painful wasp sting!”

Based on these considerations we hypothesised that there should be an observable boundary to the modulation of pain perception by expectation. We hypothesized that when individuals are presented with increasingly discrepant sensory evidence – and increasingly large prediction error – the influence of expectation on perception will decrease. To test this hypothesis we delivered pain stimulus intensities that violated cued expectations to increasing degrees and tested the effect on the resulting pain intensity rating. Our hypothesis would be supported if the influence of cued pain intensity on pain intensity rating decreased on trials with a large PE, namely, those with highly unexpected pain stimulus intensities. This would also support our hypothesis that PE has a functional role in pain perception.

In two experiments, we varied the cued intensity and the stimulus intensity of a painful stimulus across trials, and measured pain ratings. The experiments were conducted on independent samples by different experimenters; the second experiment was pre-registered (<u>osf.io/5r6z7</u>). On a given trial, PE was defined as the numerical difference between cued intensity and stimulus intensity. For example, when the cue signalled that the pain would be a 2 on a 1-10 pain scale, and the stimulus intensity was 4 on the same scale, the PE on that trial was +2 (PE is depicted on the X axis in figure 1). The outcome variable (the Y axis in figure 1) was the numerical difference between the stimulus intensity and the pain rating the participant gave on that trial, which we term subjective error, or PE_sub_. In this example, if the participant was biased by the cue, and rated the stimulus not as a 4 but slightly lower, as a 3, the PE_sub_ would be −1. Therefore, PE_sub_ measures how much pain intensity ratings were influenced by the cue on a Trial-by-Trial basis, both when pain is greater than expected - as in placebo analgesia (positive PE, on the right side of figure 1), as well when pain that is lower than expected - as in nocebo hyperalgesia (negative PE, on the left side of figure 1). In both cases, a small absolute PE_sub_ indicates pain experience close to the actual stimulus intensity, whereas a large absolute PE_sub_ suggests that the pain experience was influenced by factors other than the incoming stimulus intensity. The predicted relationship between PE and PE_sub_ are depicted schematically in Figure 1. Our hypothesis would be supported if we observe a ‘tipping point’ where pain stimulus intensity is so discrepant to expectation that the influence of expectations on perceived pain intensity decreases. We predicted that this would occur both when PE was positive and when it was negative, namely, in both the ‘placebo’ and the ‘nocebo’ conditions of the two experiments.

**Figure 1.**
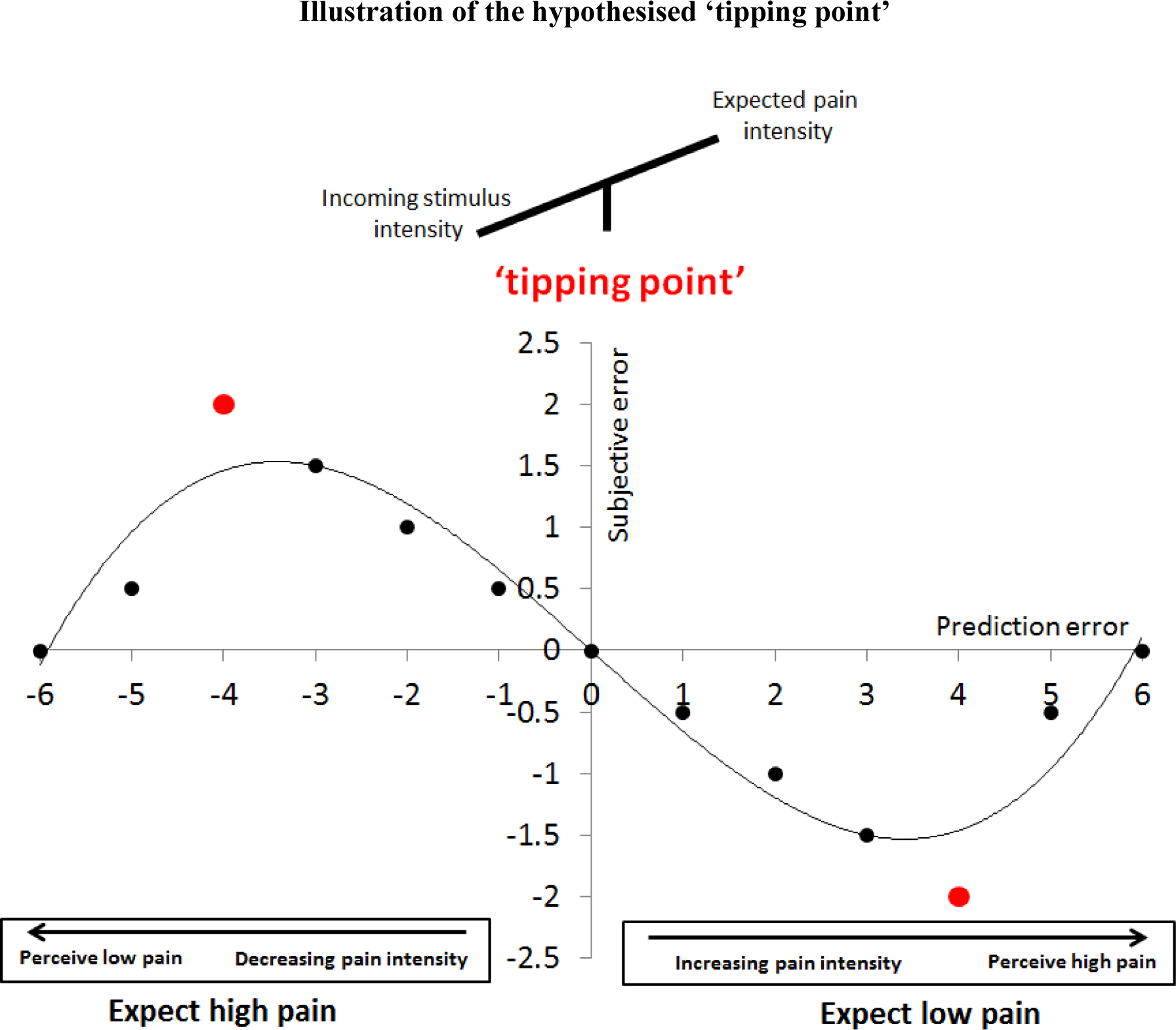
Graphical representation of the hypothesised polynomial relationship between PE and PE_sub_. As the discrepancy between cued intensity and stimulus intensity (PE) increases (from the origin towards both extremes of the X axis), the discrepancy between stimulus intensity and pain intensity rating (PE_sub_) also increases, as expectations influence pain perception. The ‘tipping point’ (red marker) is reached where stimulus intensity is so unexpected that the influence of expectations begins to decrease. Positive PE, where pain was greater than expected, is plotted on the right side of the plot. For example, a PE of +2 would reflect a cued expectation of 2 on the NPS, but an actual stimulus intensity of 4 NPS (4 NPS – 2 NPS); as this is a small PE, the cued expectation influences perception, resulting in a perception of 3 NPS and a PE_sub_ of −1 (3 NPS – 4 NPS). A PE of +5 would reflect the same cued intensity, 2 NPS, but an actual stimulus intensity of 7 NPS; as this is a large PE, beyond the perceptual ‘tipping point’, the influence of expectation on perceived pain is decreased, resulting in a perception of 6.5 NPS, and PE_sub_ is decreased to −0.5 (6.5 NPS – 7 NPS). This hypothetical relationship is also plotted for pain that is lower than expected, associated with negative PE, on the left side of the plot. Across positive and negative PE Trials, the hypothesised relationship between PE and PE_sub_ would be best expressed by a cubic polynomial.

First, we analysed the relationship between cued intensity and pain intensity ratings in Dataset1 to check that our pain cueing manipulation had the desired effect. Second, we used regression modelling to evaluate the relationship between PE and PE_sub_. A cubic polynomial model was fitted to Dataset 1 to test our hypothesis of a non-monotonic relationship between PE and PE_sub_ for both positive and negative PE. Such a non-monotonic relationship would indicate that cued intensity influenced pain intensity rating up to a certain threshold (the ‘tipping point’), reflected in an increase in PE_sub_, and that this influence decreased when stimulus intensity was highly unexpected (high absolute PE), reflected in a decrease in PE_sub_. Third, we developed a more complex model to explore additional effects in the dataset. Motivated by findings that expectations of low pain, as in placebo analgesia, are more easily extinguished and are more sensitive to sensory evidence than expectations of high pain, as in nocebo hyperalgesia^8, 18–20^, the more complex model included a quartic term to uncover any differences in the effect of expectations on pain that depended on the sign of the PE. The model also examined whether the effects of expectations changed throughout the course of the experiment, for example due to learning, habituation, sensitisation, or fatigue. In this model we also tested for individual variation in the relationship between PE and PE_sub_. Finally, to validate our conclusions, we pre-registered the experimental design and the analysis plan, and collected a second, independent Dataset 2, to which we applied the same two models.

## Results

### Manipulation check: Effect of cued intensity on pain rating

We first assessed the extent to which cued intensity and stimulus intensity influenced pain intensity ratings. Figure 2 plots averaged pain ratings from Dataset 1 as a function of stimulus intensity. The plot suggests, unsurprisingly, that pain ratings increased when stimulus intensity increased. As expected, a stimulus preceded by a level 2 cue (black) was rated as lower than the same intensity stimulus preceded by a level 8 cue (grey). We analysed this relationship with a regression model (table 1). The model explained around 58% of the variation in pain ratings. The results (table 1) indicated that stimulus intensity, cued intensity and the interaction between them were all significant predictors of pain intensity ratings. As cued intensity and stimulus intensity increased, so did pain intensity rating, but the larger beta value for the effect of stimulus intensity (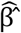=0.63) than cued intensity (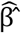=0.23) indicated that the influence of the former were greater. The significant interaction term also indicated that the effects of cued intensity increased as the stimulus intensity increased, and vice versa. These results indicate that cued intensity had the desired influence on pain ratings.

**Figure 2.**
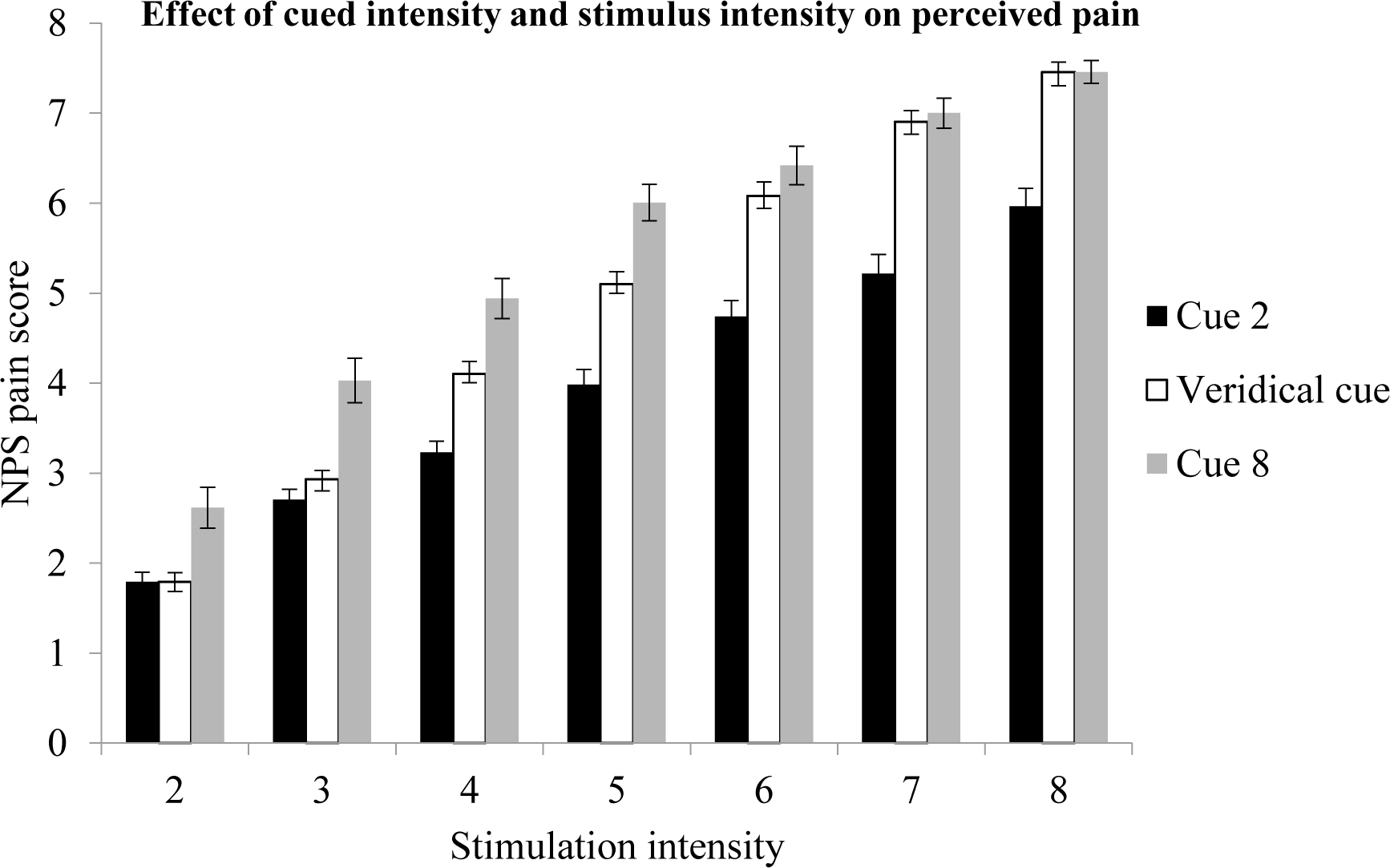
Average influence of cued intensity and stimulus intensity on NPS rating. A cue predicting low intensity pain (Cue 2) decreases the perceived intensity of a painful stimulus compared to baseline (veridical cue), and a cue predicting high intensity pain (Cue 8) increases the perceived intensity compared to baseline. Error bars represent the standard error of the mean.

**Table 1:**
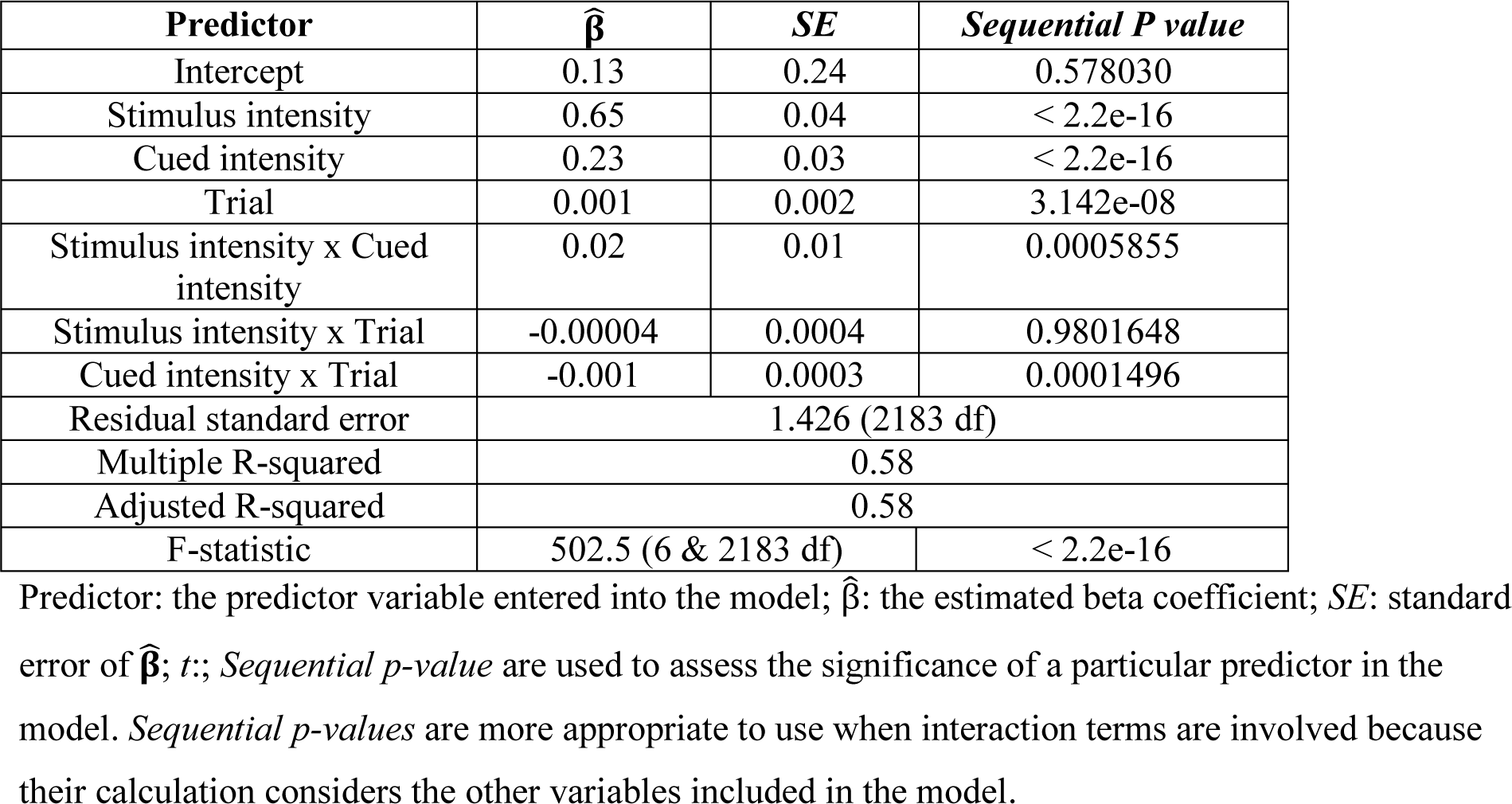
Results of the fixed effects model on Dataset 1, assessing the extent to which stimulus intensity, cued intensity and trial influenced pain intensity rating

The results also showed that trial was a significant predictor of pain rating. The significant interaction between cued intensity and trial indicated that the effect of the cues on pain ratings changed throughout the course of the experiment. The interaction of trial and stimulus intensity did not significantly predict pain rating. These results motivated us to explore the effects of Trial further, as described under ‘complex model’, below.

### Relationship between PE and PE_sub_

#### Basic model

We next tested whether the modulation of pain rating by cue was influenced by PE, the difference between cued and stimulus intensity (table 2). We predicted that when stimulus intensity is moderately different from the cued intensity, namely when PE size is moderate, PE_subj_ would be the largest, as shown around the middle of both the positive and negative X axes on figure 1. We also predicted that when stimulus intensity resembled expectation, and, more controversially, when it was very different from expectation – namely when PE was either small or large – PE_sub_ would decrease, as shown furthest from the axes origin in figure 1. A cubic polynomial describes this predicted relationship between PE and PE_sub_.

**Table 2:** Results of the basic models fitted in Dataset 1 and 2

| Polynomial model: $PE_{sub}$ | | | | | | |
| --- | --- | --- | --- | --- | --- | --- |
|  | Dataset 1 |  |  | Dataset 2 |  |  |
| Predictor | $\hat{\beta}$ | SE | Marginal P-value* | $\hat{\beta}$ | SE | Marginal P-value* |
| Intercept | -.13 | .04 | .003 | -.44 | .05 | < .0001 |
| PE | -.33 | .02 | <.0001 | -.27 | .02 | <.001 |
| $PE^2$ | -.01 | .002 | <.0001 | -.01 | .002 | <.001 |
| $PE^3$ | .002 | .001 | .0003 | .004 | .001 | <.0001 |
| AIC | 7794.981 |  |  | 7639.873 |  |  |
| BIC | 7823.44 |  |  | 7668.121 |  |  |
| Log-likelihood | -3892.491 |  |  | -3814.936 |  |  |

Predictor: the predictor variable entered into the basic model; $\hat{\beta}$ : the estimated beta coefficient; SE: standard error of $\hat{\beta}$ ; Marginal P-values are used to assess the significance of a particular predictor in the presence of no other predictors in the model.

The results from fitting the cubic polynomial fixed-effect model (henceforth, ‘basic’ model) to Dataset 1 indicate that all the terms are significant (table 2). Note that the magnitudes of the estimates for polynomial terms should not be compared directly, because they are on different scales: PE takes values between 0-6, whereas the values taken by PE^2^ and PE^3^ range between 0-36 and 0-216 respectively. Therefore, the lower magnitude of the beta estimates for the PE^2^ and, crucially, the PE^3^ terms, compared to the estimates for the PE term does not reflect the magnitude of their contribution to explained variance. Adjusted R^2^ values for the models fitted on both Datasets indicate that the basic model accounts for 31% of variance of Dataset 1, and 17% of variance for dataset 2 (table 3), suggesting that there is scope to develop a more extensive model to account for additional variance.

**Table 3:** Multiple and adjusted R^2^ values for the basic model for Datasets 1 & 2

|  | **Dataset 1** | **Dataset 2** |
| --- | --- | --- |
| **Multiple $R^2$** | 0.31 | 0.18 |
| **Adjusted $R^2$** | 0.31 | 0.17 |

The multiple $R^2$ explains the percentage of variance explained by the model. The adjusted $R^2$ penalises $R$ so that it does not automatically increase with the addition of more predictors to the model but only increases with the addition of variables that increase the explained variance by a significant amount.

Figure 3 shows the curves of the relationship between PE and PE_sub_. In Dataset 1the curve reflects the predicted cubic relationship between PE and PE_sub_: as PE increases (in either a negative or positive direction), PE_sub_ also increases, but at the higher levels of PE (most negative or most positive), PE_sub_ decreases. This mirrors the relationship we hypothesised, depicted in figure 1. Although these results support our hypothesis of a ‘tipping point’ at high size of positive PE, the ‘tipping point’ effect was subtle. Its modest magnitude brought the robustness of this relationship into question. We thus pre-registered the experimental design and the model, collected a second, independent dataset (Dataset 2), and applied the same model to it, to test whether the predicted relationship would be replicated. As can be seen in figure 3, the second Dataset more clearly reflects the cubic relationship, suggesting that the effect is robust and supporting our hypothesis of a ‘tipping point’ at high levels of absolute PE.

**Figure 3.**
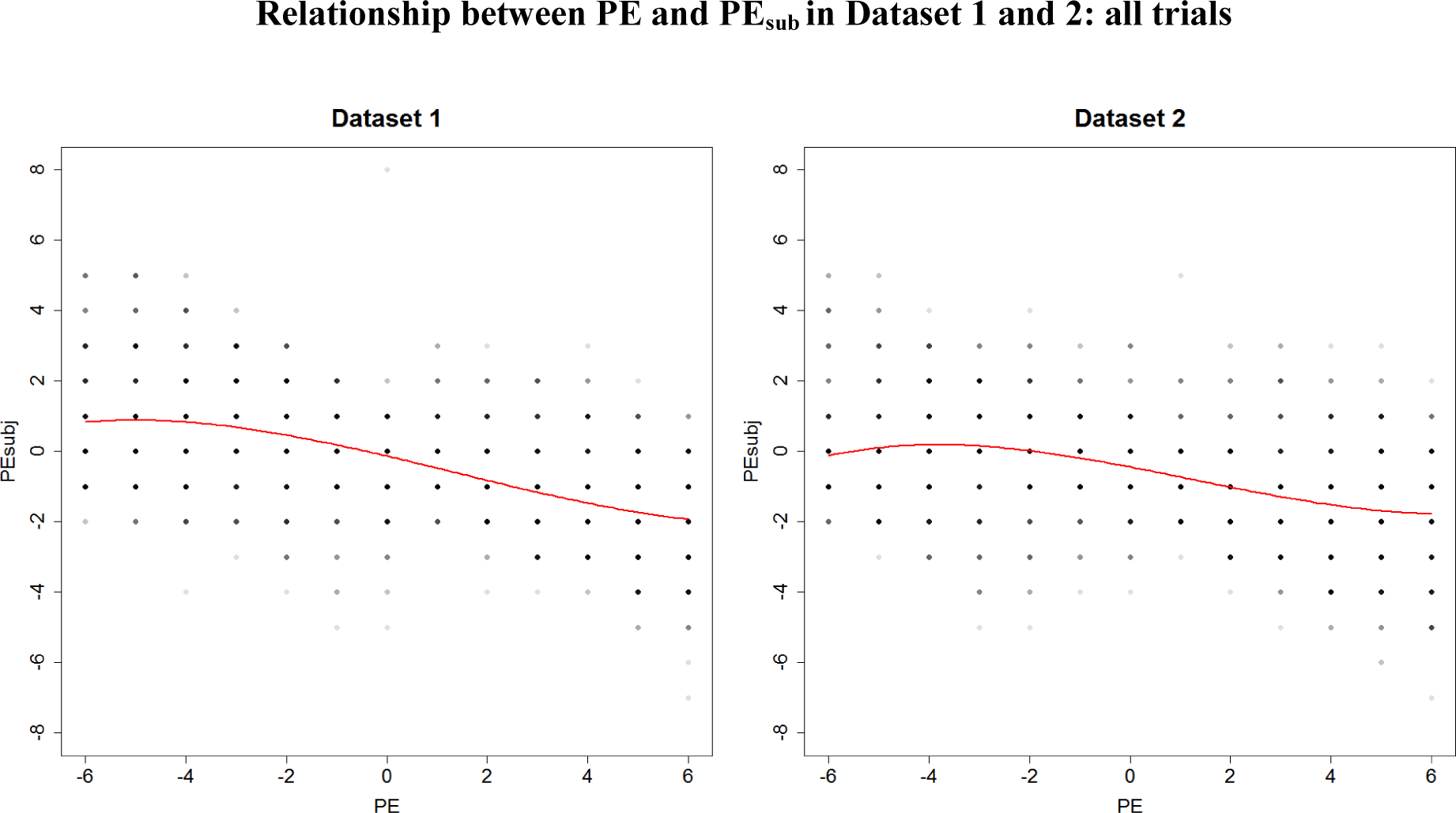
Initial fitted models per dataset, both of which show a cubic relationship between PE and PE_sub,._ This relationship is particularly clear in Dataset 2.

#### Complex model

The basic model confirmed our hypothesised cubic relationship between PE and PE_sub._ However, it is important to acknowledge that there were some key variables in the experiment which it did not account for. These were putative differences in the impact of receiving pain that is higher, compared to lower, than expected^18–21;^ potential change in pain perception or in the effect of expectation over time during the course of the experimental session; and variation between individuals in the relationship between PE and PE_sub_. To address these differences we applied a more complex model to the two Datasets. Crucially, the complex model was developed on Dataset 1, preregistered before collecting Dataset 2, and applied to Dataset 2 without any alterations.

We began by adding a 4^th^ order quartic term (PE^4^) in the model to reveal differences in the relationship between PE and PE_sub_ between the positive and negative PE conditions. The statistical motivation to include this term was explored in a preliminary analysis of Dataset 1, which indicated that the quartic term should be included alongside the cubic term (Supplementary Materials online). Supplementary Figure S1 online indicates that for some individuals in Dataset 1, e.g. participants 11, 14 and 28, a quartic polynomial is appropriate. We therefore included polynomial PE terms up to the 4^th^ order.

Theoretical models of the influence of expectation on perception refers to the instantaneous perceptual response to a sensory stimulus^10, 22^. However, it is important to note that responses to a cue could change over the course of the experimental session due to factors such as fatigue, habituation or learning ^23–28^. Furthermore, experiencing PEs could, over time, change participant’s association between the cue and the pain outcome ^15, 16, 29^. To investigate changes in the relationship between PE and PE_sub_ over time we also included Trial as a predictor, as well as its interaction with the polynomial PE terms. Henceforth, we refer to the resulting quartic polynomial model that included the effects of Trial as the ‘complex’ model. Note that Trial effects on the relationship between PE and PE_sub_ can stem from one or more of the factors mentioned above.

The relationship between PE and PE_sub_ is likely to vary between individuals. We investigated the importance of individual differences by comparing versions of the quartic polynomial model that included the effect of Trial in order to identify which random effects were needed in the model. Using the likelihood ratio test, we compared a model without random effects, with just the random intercept, and with both random intercept and random slope of the polynomial PE terms. The results verified that both the random intercept and random slope for the linear PE term should be included (Supplementary Materials online). The addition of random slopes in PE^2^, PE^3^ and PE^4^ did not offer significant improvements to the model.

Table 4 summarises the results of the complex model, fitted on Datasets 1 and 2. Both sets of results suggest that a quartic model in PE, with an effect for Trial and some interactions between Trial and PE are needed to describe the relationship between PE and PE_sub._ The presence of the quartic term indicates a difference in the relationship between PE and PE_sub_ between the positive and negative PE conditions. Figure 4 visualises the relationship between PE and PE_sub_ based on this model, at different levels of Trial, for Datasets 1 and 2.

**Figure 4.**
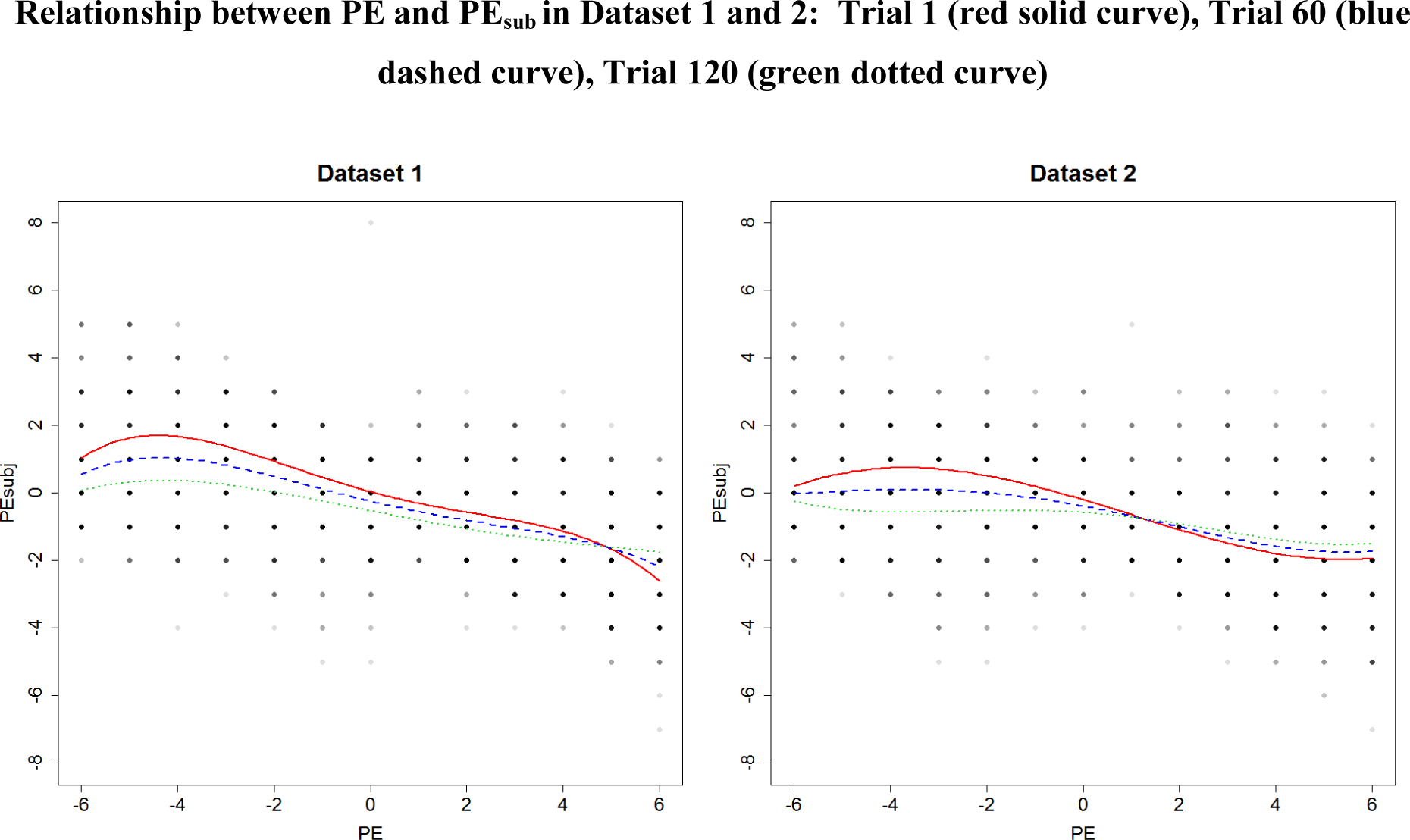
Polynomial relationship between PE and PE_sub_ per dataset; Trial 1 (red solid curve), Trial 60 (blue dashed curve), Trial 120 (green dotted curve)

**Table 4:**
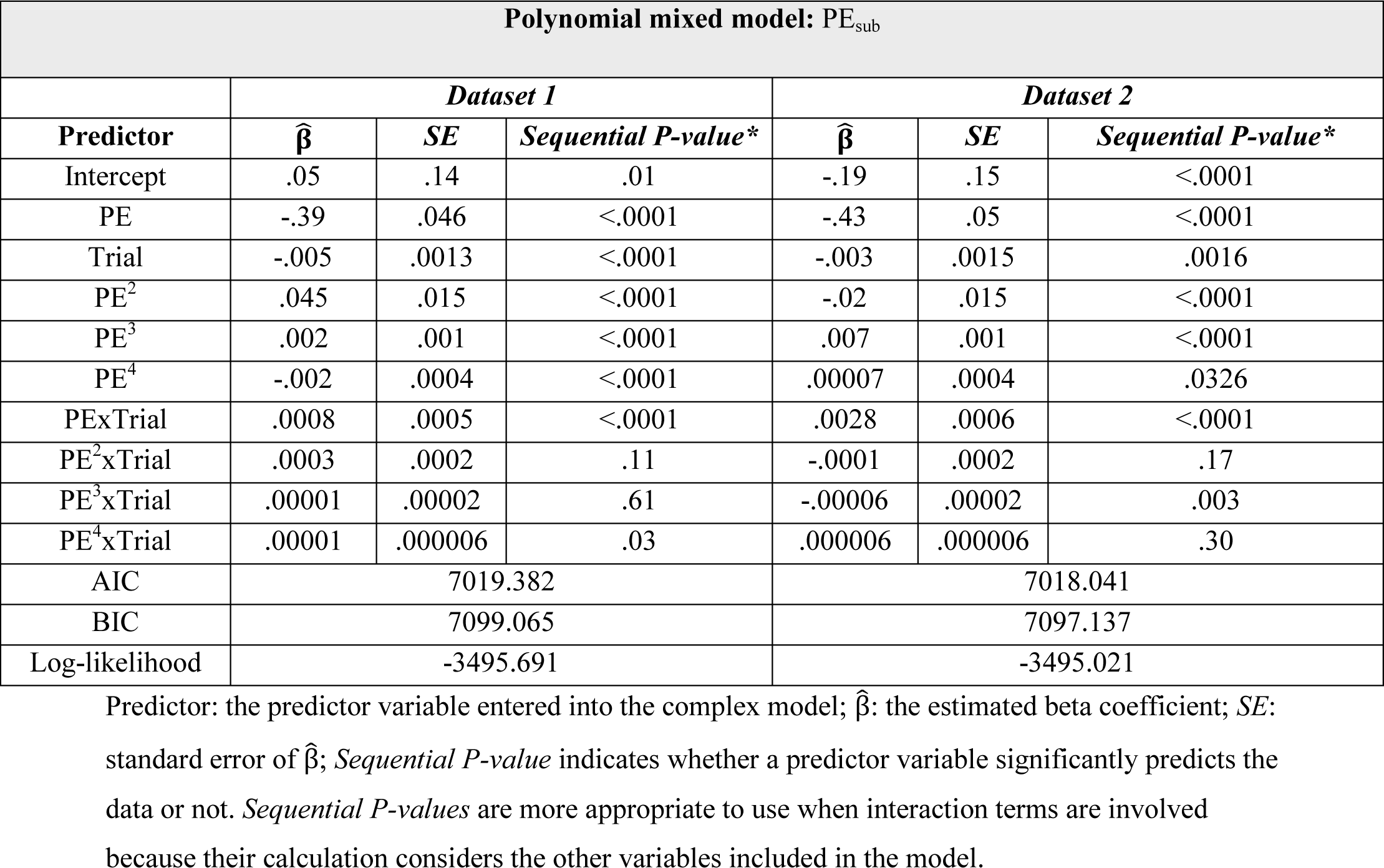
Results of the complex models fitted in Dataset 1 and 2

While the main effects as well as the interactions between PE and Trial were significant in both datasets, and in neither was the interaction between PE^2^ and Trial significant, there was no agreement across datasets as to the significance of the interactions between the higher order polynomial terms (PE^3^ and PE^4^) and Trial. This suggests that the change in the effects pain expectations had on perception during the course of the experimental session was slightly different in the two Datasets. Figure 4 reflects some subtle differences in the effect of Trial between the two datasets. In Dataset 2, the ‘tipping point’ is most pronounced at the beginning of the experimental session and becomes less pronounced over the course of the task. At the beginning of the experimental session (Trial 1, red solid curve) the curve’s polynomial cubic shape is more pronounced and clearly supports our hypothesis about the presence of a “tipping point” in response to both positive and negative PE. As participants progress through the task to Trial 60, the curve retains its earlier shape (though less pronounced). At Trial 120 (the final Trial, green dotted curve), the curve appears more linear, which suggests that that the relationship between PE and PE_sub_ weakened at the end of the experiment, indicating that the ‘tipping point’ effect decreased over the course of the task. The curve also moves closer to PE_sub_= 0, which suggests that the influence of expectation on average decreased over the task. Dataset 1 shows the same pattern of results in the negative PE condition but a slightly different effect of Trial in the positive PE condition. Here, the ‘tipping point’ is least pronounced at the beginning of the study session (Trial 1). Progressing to Trial 60 (blue dashed curve), the curve becomes more pronounced, and is most pronounced by Trial 120. These results indicate that there is some variation in how the significant ‘tipping point’ effect is expressed between the two datasets when PE is positive, namely, when pain is greater than expected.

The importance of variation between participants is evident in the comparison of marginal and conditional R^2^. The marginal R^2^ denotes the variation in the data explained only by the fixed effects of the model, whereas the conditional R^2^ denotes the variation explained by both the fixed and the random effects of the model^30^. The marginal R^2^ values indicate that the fixed effects in the complex model account for 33% (19%) of the total variation in dataset 1(2) and not surprisingly these values are similar to the R^2^ values of the basic model (which only includes fixed effects). The conditional R^2^ values indicate that the fixed and random effects account for 55% (42%) of the total variation in dataset 1(2), providing evidence in support of the complex over the basic model (table 5). The presence of a significant random intercept in the model indicates that each individual’s PE_sub_ varied significantly when PE was zero, namely, in response to veridical cues. This interpretation is supported by the visualisation of individual intercepts in Supplementary figures S1 and S2, which varies between individuals.

**Table 5:** Marginal and Conditional R^2^ values for the complex model for Datasets 1 & 2

|  | **Dataset 1** | **Dataset 2** |
| --- | --- | --- |
| **Marginal $R^2$** | 0.33 | 0.19 |
| **Conditional $R^2$** | 0.55 | 0.42 |

The marginal $R^2$ denotes the variation explained only by the fixed effects whereas the conditional $R^2$ denotes the variation explained by both fixed and random effects.

It is also possible to explore how well the model captures individuals’ behaviours, by visualising the curves for each participant in Dataset 1 (Supplementary Figure S1 online). In general, the model-fitted values (black line) capture the data well. However, for some individuals in Dataset 1, such as participants 6 and 12, the curve based on the fitted values seems to ignore the more extreme PE_sub_ values, thus producing a flatter curve. A similar pattern of results was observed for Dataset 2 (Supplementary Figure S2 online). This indicates that whilst some individuals expressed a quartic relationship between PE and PE_sub_, other participants expressed a more linear relationship.

## Discussion

We aimed to test for a ‘tipping point’ or a boundary effect on the influence of expectation on pain perception. We hypothesized a non-monotonic relationship between how discrepant the intensity of a painful stimulus is from expectation (PE), and the influence of expectation on pain perception (PE_sub_). This is the first study to systematically vary the size of pain PE and test the influence on the resulting pain perception. Specifically, we predicted that the expectation would influence on perception most strongly when stimulus intensity was moderately different from the expected intensity, when PE was moderate in size (the middle of the scale in figure 1 in either the positive or negative direction). Controversially, we also predicted that its effect would decrease when stimulus intensity was highly discrepant from the expected intensity, when PE was at its highest (furthest from the origin of figure 1 in either positive or negative direction). Our preregistered hypothesis was implemented in a cubic polynomial regression model where PE predicted PE_sub_. Across two independent datasets, the results confirmed our hypothesis. Returning to the example of a garden picnic, this suggests that a highly unexpected wasp sting would mean that the expectation of a pain-free picnic would give way to the immediate sensory data. These results extend our understanding of the relationship between expectation and perception, because they suggest that there is a limit to expectation’s influence on immediate perception.

In order to test our hypothesis about the relationship between PE and PE_sub_, where beyond a certain threshold absolute PE_sub_ decreased when absolute PE increased, we first fitted a regression model, referred to as the ‘basic’ model, which included cubic polynomial terms in PE. In Dataset 1 the result of the model supported our hypothesis. However, for positive PEs (where pain was higher than expected) the effect was small despite being significant. To further examine whether this was a true effect, we collected a second, independent Dataset 2 and repeated the same model. The basic model in Dataset 2 replicated Dataset 1, now with an equally pronounced cubic relationship between both positive and negative PEs and PE_sub_.

We also implemented a more complex model which crucially allowed us to test for three potential sources of variation in the data. First, we included a quartic term in the model to account for potential differences between the positive and negative PE condition. Second, the complex model allowed us to investigate the effect of Trial and interactions between Trial and the effect of cued intensity on pain rating. Third, the complex model allowed us to explore whether individual variation on the ‘tipping point’ was significant by testing for random as well as fixed effects. This analysis, referred to as the ‘complex’ model, too was preregistered prior to the collection of Dataset 2 (<u>osf.io/5r6z7</u>). We review findings relevant to each one of these elements below.

Our positive PE condition is comparable to a placebo manipulation (expect low pain) and our negative PE condition is comparable to a nocebo manipulation (expect high pain) ^31^. We hypothesized a quadratic relationship between PE and PE_subj_ within each of these conditions, separately when PE is positive or negative, as described in figure 1. We combined these two quadratic relationships to a single model with a cubic term in the basic model. Notably, there was no reason to hypothesise that the quadratic relationship between PE and PEsubj was of equal magnitude in the ‘placebo’ (positive PE) or ‘nocebo’ (negative PE) direction; the quartic term in the complex model allowed us to capture differences between these conditions. Previous work suggests that such differences are entirely likely. Nocebo responses are stronger, more difficult to extinguish and less sensitive to sensory evidence than placebo responses^8, 18–20^. For example, nocebo effects are equal in magnitude in response to either verbal suggestions or a conditioning procedure, whereas a conditioning procedure produces a greater placebo response than verbal suggestions alone^8^. Nocebo responses are persistent and difficult to extinguish whereas placebo responses are easier to extinguish ^18, 20^. Nocebo responses have been shown to be significantly stronger than placebo responses in a subliminal pain conditioning procedure, and this finding was related to the higher salience of nocebo cues ^19^. Placebo and nocebo responses are also associated with different neurotransmitters; placebo responses are associated with the release of dopamine and endogenous opioids, whereas nocebo responses are associated with the release of cholecystokinin^32–34^.

The main difference in result between the positive and negative condition here was that boundary effects were less consistent in the positive PE condition, in the sense that they depended on Trial in Dataset 1 but not Dataset 2. This did not happen in the negative PE ‘nocebo’ condition. It is plausible that because expectations of low pain are more easily extinguished and more sensitive to sensory evidence^18–21^, the corresponding positive PE condition was more vulnerable to an unmeasured variable than the negative PE condition. Future research on the boundary effects of expectations should investigate the correlates of Trial, such as sensitisation, habituation, fatigue and learning. The difference between the two datasets also hints at potential differences between the samples, perhaps due to individual differences that were not measured or analysed in the present work. One variable, for example, could be individual personality traits. The trajectory of placebo responses is sensitive to dispositional optimism ^4^. It is possible that by chance there was a higher proportion of optimists, or of some other unobserved individual trait, in Dataset 1 compared to Dataset 2, which influenced their learning from higher-than-expected pain. Our modelling approach afforded a quantification of boundary effects at the individual level, to enable future work to explore the predictive power of individual personality traits on the relationship of PE vs PE_sub_.

The complex model revealed that the effect of cue changed over the course of the experimental session. Indeed, changes over time could be the source of the differing average relationships between PE and PE_sub_ between certain participants (Supplementary figures S1 and S2 online). Our interest in how perception changes across the experimental session was motivated by the observation that sensitivity to painful stimulation can vary over time ^15, 16, 28, 29^. The computational mechanism driving this change still needs to be worked out. Current perspectives on the perceptual mechanism behind expectation effects ^10, 35–38^ suggest that an agent maintains multiple potential hypotheses about the cause of a sensory input, and selects the one which best represents the incoming sensory information at any given time. In our results, it is possible that through the experience they had over the course of the task, the weight participants assigned to competing hypotheses about the causes of sensory input began to shift, resulting in a change in the magnitude of boundary effects. Future work should model this process computationally to elucidate the mechanism behind it.

Our interest in individual differences was motivated by our own and others’ findings that the experience of pain is strongly affected by other psychological variables such as optimism, anxiety, suggestibility, and reward responsiveness ^1–5^. It was important to consider individual differences here especially because we previously showed that they account for variation in pain expectation effects on pain experience ^22^. The complex mixed model was a significantly better fit than the basic fixed model and this was supported by the higher R^2^ for the complex model when both fixed and random effects were considered. The presence of a significant random intercept in the complex model indicates some change in response to the painful stimulus between individuals when the pain stimulus intensity matched the cued intensity (PE=0). It is possible that some participants did not rate the stimulus intensity the same as they had rated it in the calibration session due to a time-sensitive factor such as habituation or sensitisation^23–28^.

Our results only provide behavioural evidence for a role of prediction error in pain and we do not claim to elucidate the neural mechanism of the ‘tipping point’. There are many putative neural mechanism behind the influence of expectation on pain perception ^39–42^. One pathway which could be responsible for the boundary effects of expectations was proposed within an influential review of placebo analgesia and expectation, in the context of predictive coding theory of pain perception^10^. This was namely the periaqueductal gray-rostral ventral medulla-spinal cord (PAG-RVM-SC). The neural expression of PE has been captured in the PAG, and the PAG-RVM-SC has been identified as a potential pathway for the influence of expectations on perception, where endogenous opioids in this pathway signal the influence of top-down expectations ^10^. In the context of our results, the difference between cued intensity and stimulus intensity could be calculated in the PAG. When the stimulus intensity is very different to the cued intensity and the influence of expectations is decreased, this may be expressed in altered signalling from the PAG ^10, 43–45^. Future studies could repeat this study using fMRI to test whether the non-monotonic relationship we observed between PE and PE_sub_ is visible in the PAG.

There are some limitations to this study. First, we induced expectations of both high and low pain. The experience of expecting high pain is affectively different to expecting a low pain. For example, expecting low pain may decrease anxiety, whereas expecting high pain may increase anxiety, and anxiety is known to influence the perceived intensity of pain which could interact with the effect of expectation ^46^. Expecting high pain increases attention to the stimulus intensity, and attention also modulates responses to pain ^47–50^. Future studies could repeat the study while recording anxiety and attention to pain to test whether they influence the effects of pain expectations reported here. Second, we did not explicitly record participant’s expectations. This was intended to avoid interfering with the effect of cue by arousing suspicion in participants that the cues were not veridical, and is a typical method used in pain expectation studies^8, 51–53^. We instead inferred expectations from the significant effect of cued intensity on pain intensity rating (table 1). A more serious limitation is that by varying stimulus intensity here, in order to elicit different levels of PE, we also varied its salience. A higher stimulus intensity is likely to be more salient, and thus have more importance assigned to it which could increase its influence on perceived pain ^54^. It would be useful to replicate this study by varying pain intensity cues whilst maintaining a constant level of stimulus intensity, to eliminate the possibility that pain intensity, and, by reference, pain salience explained results that were attributed here to PE. Finally, in contemporary theories of perception, the influence of expectation on perceived pain is also modulated by the certainty (the inverse variability) of the pain stimulus intensity ^10, 35, 37, 55, 56^. A potential change in certainty of the cued expectation over the course of the task could relate to the different effects of Trial observed in the positive PE condition between Dataset 1 and Dataset 2.

To conclude, we explored boundary effects of expectation in pain perception, and validated our results in two independently collected datasets, the second of which was preregistered (<u>osf.io/5r6z7</u>). We show that when pain is very different to what was expected, perception moves closer to the pain stimulus intensity, especially when pain is much lower than expected. Pain perception is bewilderingly variable and pain is an important subjective experience which affects all individuals to varying degrees ^57^. Chronic pain in particular is a prevalent debilitating experience with great personal and socioeconomic costs. The economic cost of chronic pain is greater than most other conditions, and it causes suffering and significantly reduces quality of life, linked to issues in mental health, sleep, physical and cognitive functioning ^58^. Treatment for chronic pain is often ineffective and associated with undesirable side-effects ^59–62^. There is a clear need for more precise models of pain perception to inform treatment, particularly to inform non-drug interventions. Our results provide insight into the influence of the relationship between prior expectation and sensory evidence on pain perception in real time, and bring us closer to a quantitative mathematical model of pain. Furthermore, in the clinic, it is usual to give reassurance about a painful or unpleasant experience. Our results indicate that reassurance that is completely discrepant with ensuing painful or unpleasant events may not always be useful for a patient.

## Methods

### Participants

For both Dataset 1 and 2, participants aged 18-35 were recruited via university advertisements. Participants received £15 compensation. Participants had normal or corrected-to-normal vision. They had no history of neurological or psychiatric conditions, had not taken analgesics on the day of the experiment, and did not have a history of chronic pain. Ethical approval was granted by the University of Manchester, where the study took place. All methods were carried out in accordance with the Code of Ethics of the World Medical Association (Declaration of Helsinki). Informed consent was obtained from all subjects. For Dataset 1, 31 participants aged 18-35 (19 females, mean age 23 years), and for Dataset 2, 30 participants (15 females, mean age 21 years) were recruited into the study. For Dataset 1 we determined an appropriate sample size of 30 subjects by examining previous studies investigating the effect of expectation on electrical pain and more subtle effects such as the effect of certainty or subliminally presented cue on pain intensity rating, which typically recruit 15-30 participants (Brown, Seymour, El-Deredy, & Jones, 2008; Colloca & Benedetti, 2006; Jensen, Kaptchuk, Kirsch, Raicek, Lindstrom, Berna, Gollub, Ingvar, & Kong, 2012). Sample size for Dataset 2 was based on that of Dataset 1. The two studies only differed in terms of the experimenter collecting the data, and the room the data were collected in, which were nonetheless similar in shape, size and light level. They took place approximately 9 months apart.

### Apparatus

Visual stimuli were presented on a desktop computer screen one metre away from the participant. Painful stimuli were electrical pulses delivered via a concentric electrode by a constant current stimulator (Digitimer DS5 2000, Digitimer Ltd., Welwyn Garden City, UK). The pulse width of the electrical stimulation was 5 milliseconds. All stimuli were controlled through a Matlab platform (Mathworks) which interfaced with the pain stimulator via a digital-to-analogue convertor (Multifunction I/O device, National instruments, Measurement House, Berkshire, UK). Participants submitted their intensity ratings of the pain using a keypad.

### Procedure

Upon arrival to the lab, participants were briefed by the experimenter, who introduced the study as a test of pain perception. After providing consent, participants washed both hands with soap and water and the concentric electrode was attached to the back of the participant’s hand where it remained for the remainder of the session.

Participants first underwent a pain calibration procedure on their left hand to determine their response to increasing electrical stimulus intensities. The first stimulus was at a low intensity which is below the threshold for pain perception in most people. The stimulus intensity increased in a ramping procedure up to a maximum of five volts. We used a 0-10 Numerical Pain Scale (NPS) to measure the pain intensity rating, where a pain intensity rating of NPS 2 was when the stimulus became “just painful”, NPS 5 was “medium pain”, and NPS 8 was at the point where stimulus was “just tolerable”, replicating previous research ^65^. We repeated this procedure three times and computed the average stimulus intensities over these three repetitions corresponding to NPSs 2, 3, 4, 5, 6, 7 and 8. Participants then underwent a pre-experiment test procedure: stimulus intensities corresponding to their pain intensity ratings NPS 2 to 8 were delivered in a pseudo-randomised order four times and participants were instructed to identify the intensity of each pulse. Participants had to correctly identify 75% of stimulus intensities to continue to the main experiment. If they did not achieve this in the test procedure, the intensities were adjusted (intensity was increased if participants rated the stimulus intensity as lower than in the pain calibration procedure, and vice versa), and the test repeated until participants correctly identified 75% of stimulus intensities.

In the main experiment, participants were instructed that the cue predicted the stimulus intensity on each Trial. The cue was a number on the computer screen which depicted the intensity of the upcoming stimulus (Figure 5), and then a stimulus intensity was delivered which either corresponded to the cued intensity or violated it at varying levels, in a partially reinforced cueing procedure. The number of trials in each condition is detailed in Table 6. Most of the NPS 2 (“just painful”) and NPS 8 (“highest tolerable pain”) cues were followed by unexpected stimulus intensity (50% of all trials); all other pain cues were veridical, i.e. where the cued intensity matched the stimulus intensity (50% of Trials). The veridical trials reinforced participant’s belief in the validity of the cues.

**Figure 5.**
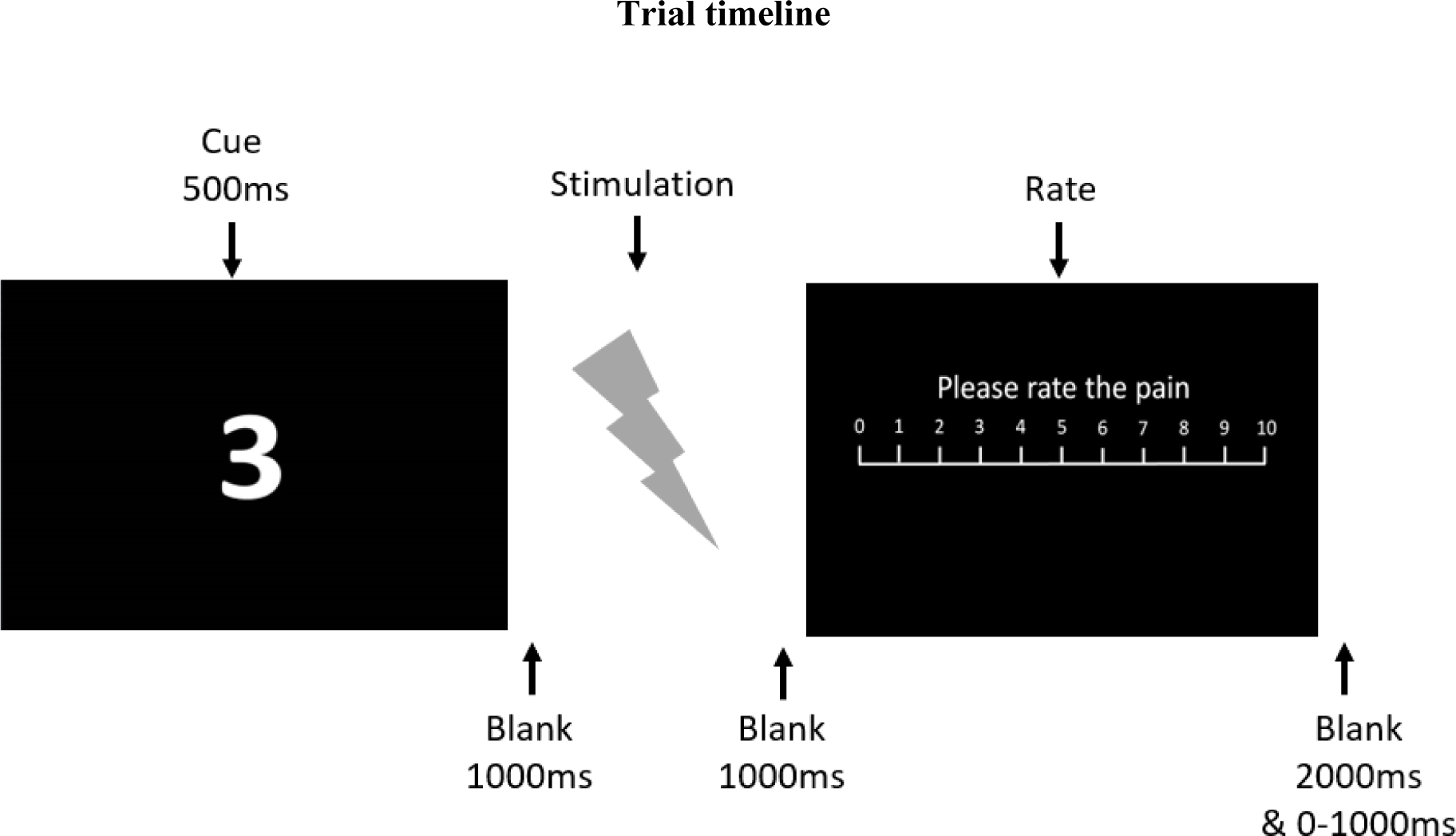
Trial timeline. After viewing a fixation cross, participants viewed a number from 2 to 8 which depicted the cued intensity for that Trial. After a blank screen, participants received the stimulus, followed by another blank screen. A rating screen was presented which prompted participants to rate the pain on a NPS scale.

**Table 6:** A summary of all Trials

| **Cued intensity** | **Stimulus intensity** | **PE** | **Number of Trials** |
| --- | --- | --- | --- |
| 2 | 2 | 0 | 5 |
| 2 | 3 | 1 | 5 |
| 2 | 4 | 2 | 5 |
| 2 | 5 | 3 | 5 |
| 2 | 6 | 4 | 5 |
| 2 | 7 | 5 | 5 |
| 2 | 8 | 6 | 5 |
| 3 | 3 | 0 | 10 |
| 4 | 4 | 0 | 10 |
| 5 | 5 | 0 | 10 |
| 6 | 6 | 0 | 10 |
| 7 | 7 | 0 | 10 |
| 8 | 8 | 0 | 5 |
| 8 | 7 | -1 | 5 |
| 8 | 6 | -2 | 5 |
| 8 | 5 | -3 | 5 |
| 8 | 4 | -4 | 5 |
| 8 | 3 | -5 | 5 |
| 8 | 2 | -6 | 5 |

Cued intensity: the level of pain stimulus intensity communicated by the visual numerical cue prior to the painful stimulus; stimulus intensity: the true intensity of the stimulus, as rated in the pain calibration procedure; PE: prediction error (stimulus intensity – cued intensity); number of trials: number of trials for that condition in the experimental session.

The cue depicted a cued intensity of NPS 2, 3, 4, 5, 6, 7 or 8 (figure 5 and table 6 for a summary of all trials in the study). In trials where PE was greater than 0, after presentation of the cued intensities 2 or 8, participants could actually receive any of the 2, 3, 4, 5, 6, 7 or 8 of stimulus intensity with the same probability. Thus a range of PEs could be elicited. Participants were instructed to rate the intensity of the stimulus and were not informed that the cues were discrepant. Trials were randomised across participants.

On each experimental trial, participants viewed a fixation cross, a cue, and then a blank screen. The stimulus was delivered, and a screen was presented which prompted participants to numerically rate their perceived pain intensity on a 0-10 NPS using a keypad. There was no time limit on this response. See figure 5.

### Data analysis

#### Manipulation check: Effect of cued intensity on pain rating

In this first model we assessed the extent to which cued intensity and stimulus intensity influenced pain intensity rating using a fixed effects model (table 1), which included the main effects for stimulus intensity, cued intensity and Trial, along with all two-way interactions between the main effects.

#### Relationship between PE and PE_sub_

##### Basic Model

We aimed to test whether PE_sub_ could be expressed as a function of PE. We analysed pain intensity ratings to stimuli which had a cued intensity of NPS 2 (low pain) or NPS 8 (high pain). PE was calculated as the numerical difference between the cued intensity and the stimulus intensity on a given trial, and so ranged from −6 (cue NPS 8 pain, deliver NPS 2 pain) to +6 (cue NPS 2 pain, deliver NPS 8 pain). This provided a measure of how discrepant the stimulus intensity was compared with expectation. PE_sub_ was calculated as the numerical difference between the stimulus intensity and the pain intensity rating on a given Trial, and so ranged, in principle, from −8 (stimulus NPS 8 pain, rate NPS 0 pain) to +8 (stimulus delivered NPS 2 pain, rated NPS 10 pain). In practice, PE_sub_ ranged from −7 to 8. This provided a measure of how far expectations shifted the pain intensity rating away from the stimulus intensity. We predicted a non-monotonic relationship between PE and PE_sub_ in line with our hypothesis of a boundary on the effect of expectation on pain perception.

The following cubic polynomial model was initially fitted to analyse the relationship between PE and PE_sub_:

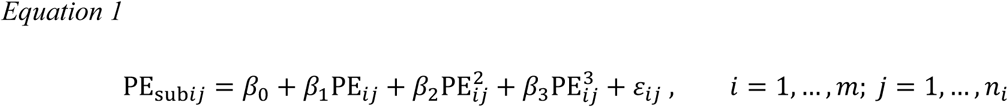

where PE*_subij_* corresponds to PE_sub_ for the *j*^th^ observation of the *i^th^* participant, PE_*ij*_ is PE for the *j^th^* observation of the *i^th^* participant, *m* is the number of participants in study and *n*_*i*_ is the number of Trials for the *i^th^* participant. We also assume that the errors ε_*i*_ = (ε_*i*1_, …, ε_*in_i_*_)^T^ ∼ *N*_*ni*_(0, σ^2^*I*_*n_i_*_), with unknown variance σ^2^,

Akaike’s Information Criteria (AIC) and Bayesian Information Criteria (BIC) ^66^values were calculated for each model to allow for model comparison.

##### Complex model

To investigate several other potential predictors, which could influence the cubic relationship between PE and PE_sub_ and account for the more subtle relationship seen in Dataset 1 (figure 3), we implemented a more complex model. This model included a quartic term, which could reveal whether the effect of pain expectations varied as a function of the direction of the difference between cued pain intensity and stimulus intensity. The quartic term allowed the ‘tipping point’ effect to have a different magnitude in the conditions with negative PE (pain lower than expected) and the positive PE (pain higher than expected).

The complex model includes polynomial terms in PE up to 4*^th^* order, a linear effect for Trial and all two-way interactions between the PE polynomial terms and Trial. The model was a mixed-effects model so it also allowed us to account for individual variation between subjects. Because including these effects resulted in a complex model, the second dataset and accompanying analysis were preregistered (<u>osf.io/5r6z7</u>).

The following linear mixed effects model with polynomial PE effects was fitted to analyse the relationship between PE and PE_sub_, as shown in equation (2):

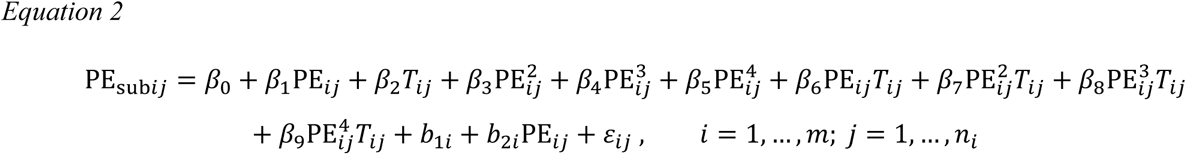

Notation here is similar to those used in equation 1; additionally, we denote *T*_*ij*_ as the Trial number for the *j^th^* observation of the *i^th^* participant, *b*_1*i*_ as the random intercept, and *b*_2*i*_ as the random slope for PE. Interactions with Trial can be interpreted as changes in the overall relationship between PE and PE_sub_ in different Trials. To see the effect of Trial more clearly, we can rearrange the model above in the following way, as shown in equation (3):

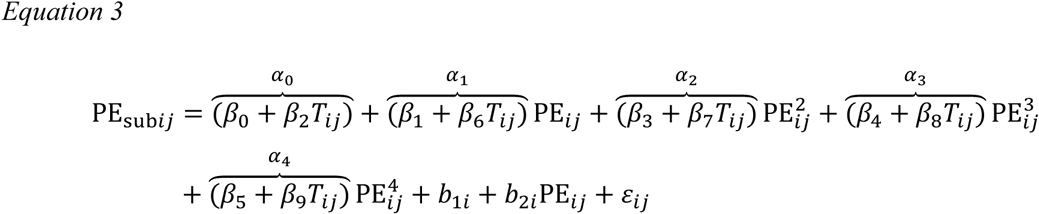

This shows that as the Trial number (*T*_*ij*_) increases, the betas of the polynomial PE terms *α*_0_, *α*_1_, *α*_2_, *α*_3_, *α*_4_ increase (decrease) if *β*_2_, *β*_6_, *β*_7_, *β*_8_, *β*_9_ are positive (negative). The above formulation also indicates that the random intercept *b*_1*i*_ accounts for differences in the intercept *α*_0_ (this is the value of PE_sub_ when PE=0) for each participant, whereas the random slope *b*_2*i*_ allows a different linear effect of PE, i.e. *α*_1_ + *b*_2*i*_, for each participant. The following assumptions are made about the random intercept and random slope for PE, as well as the measurement errors:

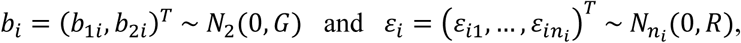

where *G* and *R* are unknown covariance matrices.

AIC and BIC values were calculated for each model to allow for model comparison.

## Acknowledgements

We would like to thank M. Parker, S. Chobert and N. Begum for their valuable contributions to this paper. This work was supported by a studentship grant from the Medical Research Council, UK. WeD acknowledges the support of CONICYT, Chile, Basal project FB0008 and FONDECYT project 1161378.

## Author Contributions

EJH: study conception, data collection, data analysis, paper writing. CC: data analysis, paper writing. WED: study conception, paper writing. AKJ: study conception, paper writing. DT: study conception, paper writing.

## Additional information

The aforementioned sources of support did not have a role in study design, collection, analysis or interpretation of data, writing of the report, or the decision to submit the article for publication. We declare no competing interests.

## Data availability

The datasets generated during and/or analysed during the current study are available from the corresponding author on reasonable request.

