## Supplementary materials for "Boundary effects of expectation in human pain perception"

Before fitting the complex model, we carried out a preliminary analysis to assess which fixed effects to include in the model, to determine the "saturated model". After visual inspection of figure 6, we checked to check whether a quartic term in PE would also be useful in describing its relationship to $PE_{sub}$, as some of the participants in Dataset 1 exhibited a quartic curve. We formally tested the significance of the quartic term by comparing the cubic PE model (basic model) with a quartic PE model through a hypothesis test. The F-test confirmed that the quartic term was significant (p-value <0.01). Adding the Trial and interaction terms in both the basic and quartic polynomial models and repeating the test further verified the significance of the quartic term (p-value <0.01).

Based on the saturated model, we then continued our preliminary analysis by exploring which random effects we needed to include in the model. Supplementary Figure S1 suggested that perhaps both a random intercept and random slope in PE would be necessary. This is because both the intercept and the linearity for each curve seem to vary across participants; compare for example participants 13 and 17. A likelihood ratio test was used to compare the saturated models without any random effects, with just random intercept and with both a random intercept and random slope, and the results confirmed that the latter model was the best (p-value <0.01).

#### The relationship between PE and $PE_{sub}$ for each subject in Dataset 1

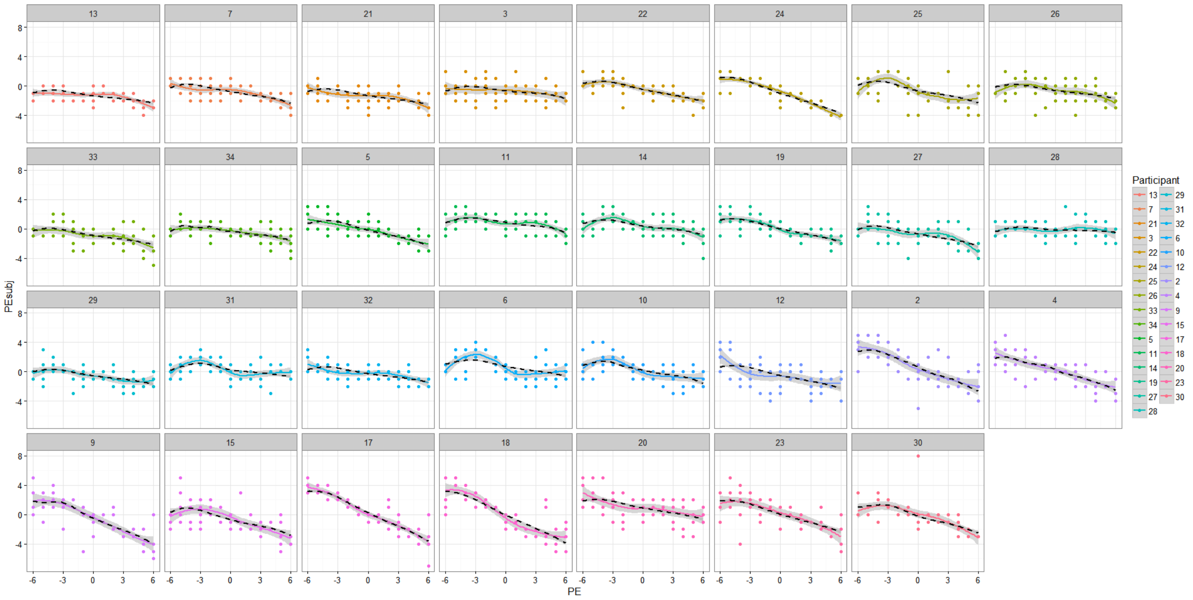


***Supplementary Figure S1:*** *Smooth trajectories illustrating the relationship between PE and $PE_{sub}$ for each subject in Dataset 1, based on the fitted values (black dashed) and based on the data (solid curves)*

#### The relationship between PE and $PE_{sub}$ for each subject in Dataset 2

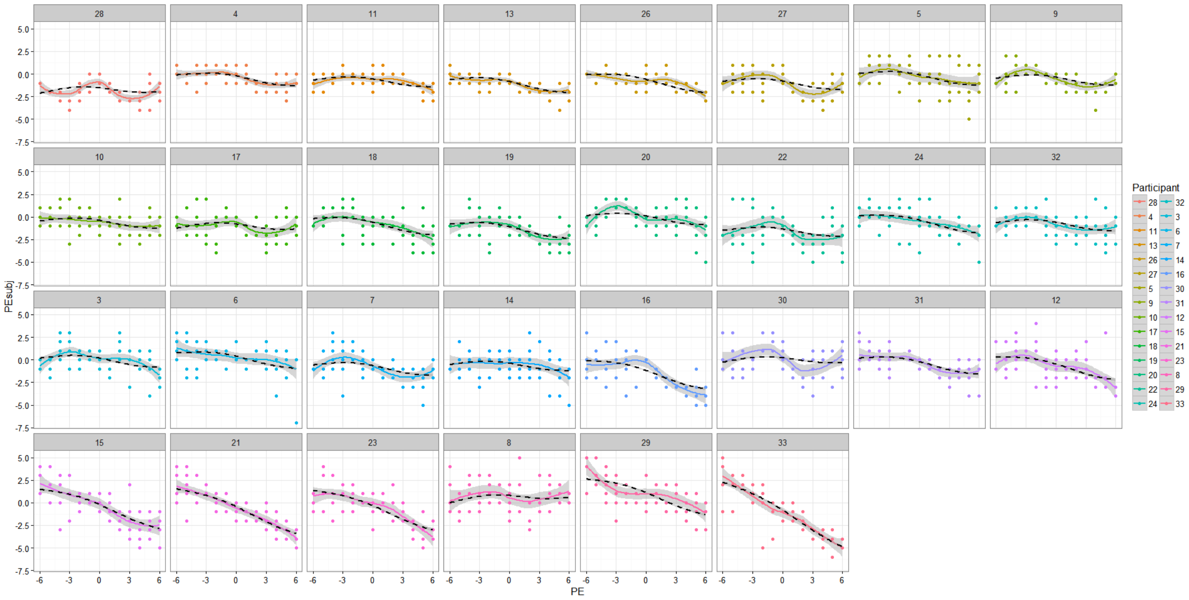

***Supplementary Figure S2:*** *Smooth trajectories illustrating the relationship between PE and $PE_{sub}$ for each subject in dataset 2, based on the fitted values (black dashed) and based on the data (solid curves)*

### Diagnostic plots for the quartic polynomial model for Dataset 1 and Dataset 2

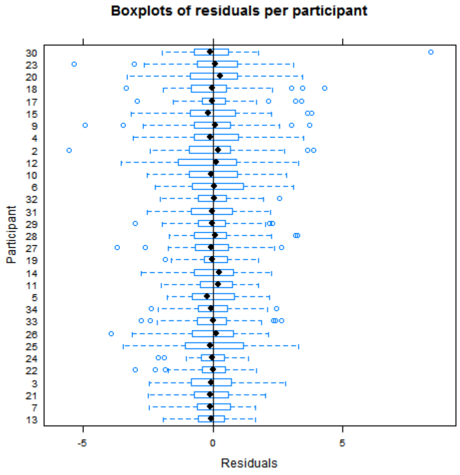


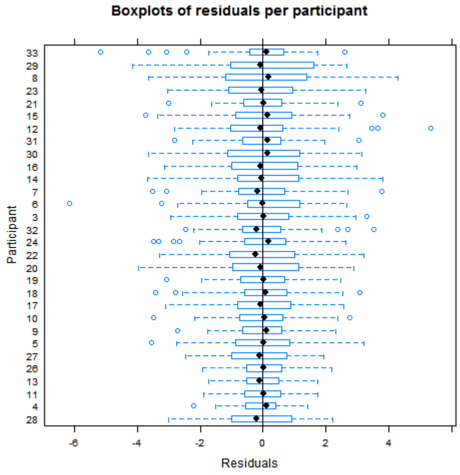


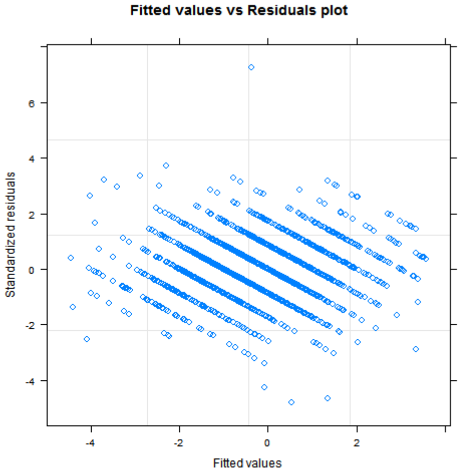


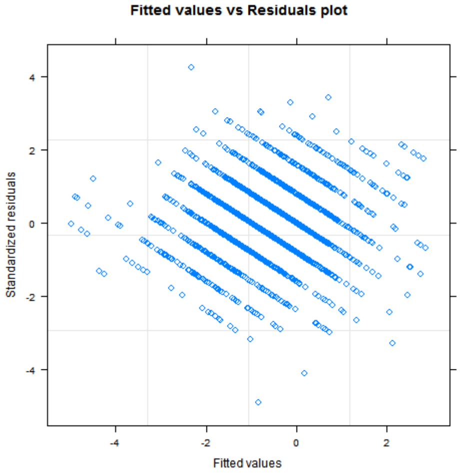


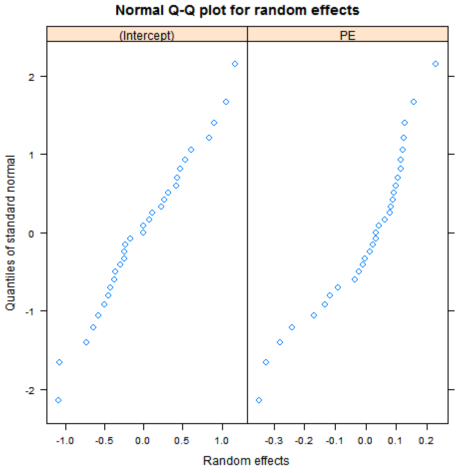


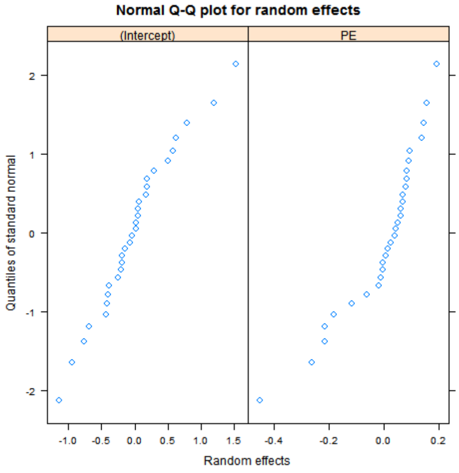

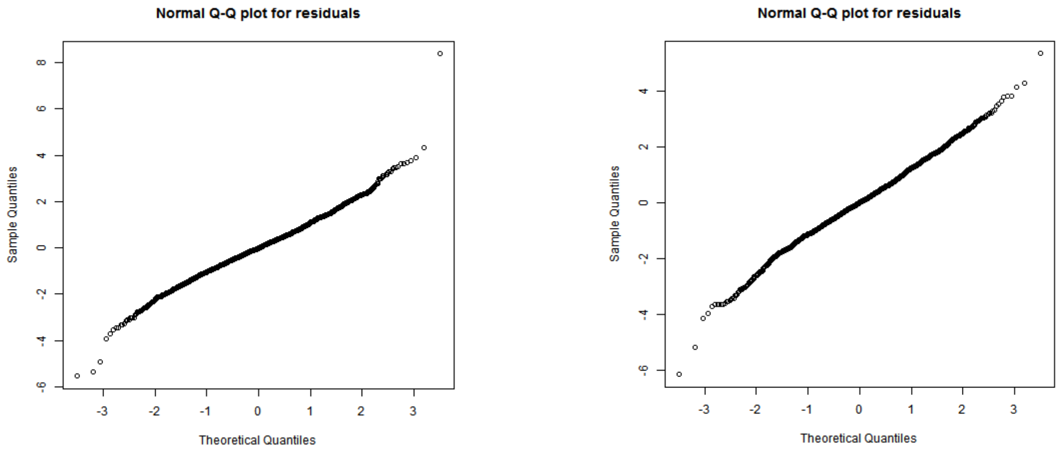


***Supplementary Figure S3:*** *Diagnostic plots for the quartic polynomial model for Dataset 1 (left column) and Dataset 2 (right column). The boxplots show the residuals are centered around 0; the plot of fitted values vs residuals shows the residuals are spread around 0 evenly (no distinguishing pattern); the qq-plots show that the Normality assumption for both random effects and residuals is reasonable.*

The results of the complex model (table 2) show that the estimates of PE, $PE^3$ and Trial are similar across datasets in terms of their sign and magnitude. This is important because these estimates reflect two key features of the relationship between PE and $PE_{sub,}$. First, at the beginning of the study (i.e. when Trial number is small), the effect of the polynomial terms dominates that of the interaction terms (because the size of the main effects coefficients is larger). Therefore, the negative sign (and size) of the linear PE term and the positive sign (and size) of the cubic PE term, ensure the tipping point for negative PE appears where it should be (as hypothesised in figure 1). Second, as Trial number increases, the negative sign of the Trial term indicates that the value attained by $PE_{sub}$ corresponding to PE = 0 shifts downwards over the course of the task. The size of the Trial term also determines how fast this shift occurs as Trial number increases. This is better visualised in Figure 3 which shows that for Trial 120 (green curve), the point at which the curve intersects the $PE_{sub}$ axis is lower compared to Trial 1 (red curve) or Trial 60 (blue curve).
